# Personalized vagino-cervical microbiome dynamics after oral probiotics

**DOI:** 10.1101/2020.06.16.155929

**Authors:** Chen Chen, Lilan Hao, Liu Tian, Liju Song, Xiaowei Zhang, Zhuye Jie, Xin Tong, Liang Xiao, Tao Zhang, Xin Jin, Xun Xu, Huanming Yang, Jian Wang, Karsten Kristiansen, Huijue Jia

**Author notes:** These authors contributed equally to this work (Chen Chen, Lilan Hao). To whom correspondence should be addressed: H.J., C.C. or.

## Abstract

The vaginal microbiota is presumably much simpler than the gut microbiome, and oral probiotics appear as a promising means to modulate its homeostasis in the general population. Here, 60 women were followed for over a year before, during and after a probiotic containing *Lactobacillus rhamnosus* GR-1 and *L. reuteri* RC-14. Shotgun metagenomic data of 1334 samples from multiple body sites did not support colonization of the probiotics to the vagino-cervical microbiome, yet the microbiome was stable in those dominated by Lactobacilli and some individuals have likely benefited from this medication-free intervention.

## INTRODUCTION

Lactobacilli have long been defined as the keystone species of the healthy vaginal microbiota. Lactic acid, hydrogen peroxide, biosurfactants and bacteriocins produced by these microorganisms help maintain the balance of vaginal microenvironment and wards off pathogens. A non-Lactobacillus-dominated microbial community has also been reported in women without symptoms of vaginosis, and is characterized by strictly anaerobic bacteria, such as *Gardnerella, Atopobium, Prevotella* and *Peptoniphilus*,which leads to significant increase in the risk of adverse conditions, including bacterial vaginosis (BV), preterm birth, urinary infections, human immunodeficiency virus (HIV), human papillomavirus (HPV) and other sexually transmitted infections (STIs)^1,2,3,4,5,6,7^.

Besides fecal transplant and dietary modulation, probiotics have become a major trend for improving gut microbiome health. E.g. for the gastrointestinal tract, gut-brain axis^8,9,10,11,12^. However, just as the vagino-cervical microbiome has received less attention as the gut microbiome, strategies for modulating the vagino-cervical microbiome is also relatively under studied.

*Lactobacillus rhamnosus* GR-1 and *L. reuteri* RC-14 are well-characterized strains as supplementations in female orally consumed probiotic products. The strains have also been reported to relieve colitis and osteoporosis in animal models^13,14^. However, evidence of their oral administration efficacy in the prevention and treatment of vaginal infection conditions, such as BV, HIV, HPV, Group B Streptococcus (GBS), remains highly debated ^15,16,17,18,19^. Moreover, the route of oral administrated probiotics to the vagina and their colonization in the multi-site of the human commensal microbiota remains largely unexplored.

Here, we conducted a longitudinal study of 60 women to explored the effect of prolonged probiotics consumption on the vagino-cervical microbiome. To investigate the dynamic alternation of muti-site microbiota after taking the live probiotic capsules, the tongue coat, buccal mucosal and fecal microbiome composition were also analyzed.

## RESULTS

### Demographic characteristics of the cohort

In our cohort, 60 healthy women were recruited (median age 31, 95% confidence interval (CI) 30-34; Supplementary Table 1). Samples were initially collected 300 days before the intervention phase. The relations of time points (before: B2, B1, B0; during: O1-O5; after: W1, W2), quantity of capsules and menstrual cycle were showed (Figure 1, Supplementary Table 2). Vagino-cervical samples were collected at all the time points. Other multi-site samples including buccal mucosa samples, tongue coat samples and fecal samples were collected at eight of the time points (B0-W2). All the samples were self-collected referring to a self-collection protocol, and performed the metagenomic analysis with shotgun sequencing data (Supplementary Figure 1). The microbial reads were extracted by filtering the human reads and subsequently used for taxonomic profiling of the microbiome (Supplementary Table 3).

**Fig. 1.**
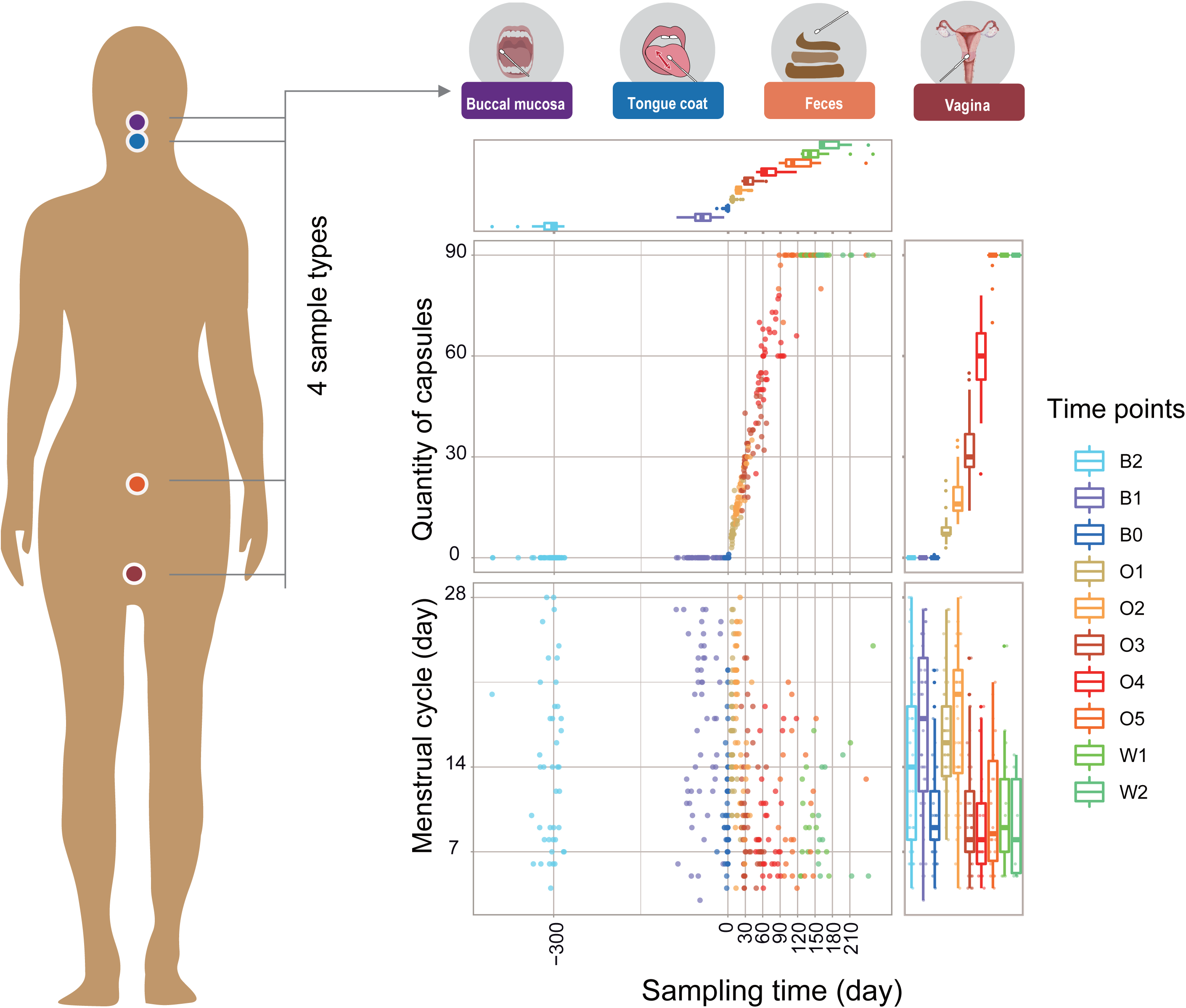
Sampling strategy of the cohort. We followed 60 healthy women for over one years each. The samples were classified into 10 time points: baseline (B2, B1, B0), during intervention period (O1-O5) and at the end of the intervention (W1, W2), according to the sampling time, quantity of capsules and menstrual cycle.

### Lack of oral probiotic colonization in the vagino-cervical microbiome

Our data showed both two probiotics were hardly present in all the body sites even during intervention period (Supplementary Figure 2a, 2b).The exception was *L. rhamnosus* GR-1 in fecal samples, which showed a weak colonization in the time-point O4 compared to the baseline (*P* = 0.01 but *q* > 0.05, Supplementary Figure 2b). Likewise, almost no change in the Shannon diversity index and Bray-Curtis dissimilarity were found between baseline and the probiotics period (Supplementary Figure 2c, 2d). We also collected the vaginal pH accompanying the sampling, no significant differences of vaginal pH were detected in all the time points (*P* = 0.87, Supplementary Figure 2e). Together, this probiotics supplementation may be limited colonization in vaginal or oral sites.

### A stable vagino-cervical microbiome is resilient against Lactobacilli intake

A previous study of the oral probiotics in individuals with BV was preceded by the antibiotic metronidazole treatment^20^, it is not clear in a more general, subclinical setting, whether the probiotic strains could really be recommended for anyone with a slight discomfort or who tested positive for potential pathogens. Compared to metagenomic data from the previous year, we classified the subjects into two groups: dysbiosis and stable, using the Bray-Curtis dissimilarity index (defined as the median Bray-Curtis dissimilarity between B2/B1 to B0) (Figure 2a). As expected, individuals of the stable group were dominated by *Lactobacillus* genera and displayed persistently lower Bray-Curtis distances, pH, Shannon alpha diversity over time compared to that of individuals in dysbiosis group (Figure 2b-2e). Thus, exogenous probiotics bacteria may be limited in impacting the vagino-cervical microbiome in stable group. However, there was still limited efficacy of probiotics in dysbiosis group (Figure2c, 2d). Of note, fecal microbiome of women in the dysbiosis group were also detected a less diverse but changed markedly compare to stable group (Supplementary Figure 3).

**Fig. 2.**
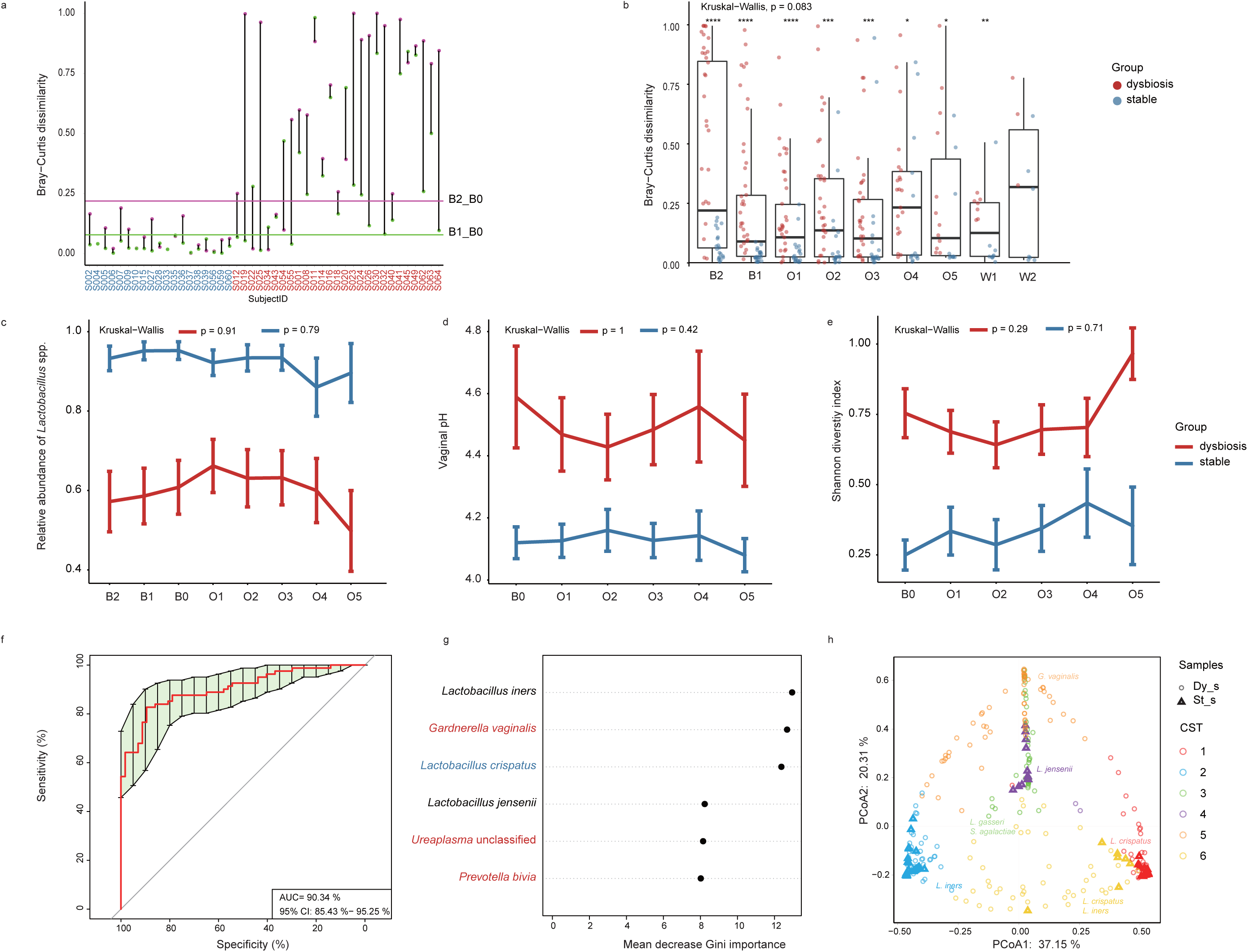
The vagino-cervical microbiome characteristics in dysbiosis and stable groups. **a, b, c, d, e**. The 46 subjects who had complete baseline time points (B2, B1, B0) were classified into two groups: dysbiosis (red) and stable (blue). **a**. Group the subjects according to the Bray-Curtis dissimilarity. Purple dots, distance between B2 and B0; green dots, distance between B1 to B0. Stable group,both dots in subjects were lower than their corresponding median Bray-Curtis distance (purple line: B2-B0; green line: B1-B0). Others were classified into dysbiosis group. **b**. The Bray-Curtis distance at each time point relative to B0. Boxplots show median and lower/upper quartiles; whiskers show inner fences. Wilcoxon ranked sum test was used to conduct comparisons between two groups in each time point, an asterisk denotes q <0.05, two asterisks denote q <0.01, three asterisks denote q <0.001, four asterisks denote q <0.0001. The Relative abundance of *Lactobacillus* spp. (**c**), vaginal pH (**d**), and Shannon diversity index (**e**) were compared between two groups. Kruskal-Wallis test was used to conduct temporal dynamics comparisons within groups. **f**. Microbiome-based discrimination between dysbiosis and stable groups. Receiver operating characteristic curve (ROC) according to 138 baseline samples (B2, B1, B0) from 27 dysbiosis subjects and 19 stable subjects calculated by cross-validated random forest models. Area under ROC (AUC) and the 95% confidence intervals are also shown. **g**. 6 species with most weight to discriminate Dy_s and St_s were selected by the models. The color of each species indicates its enrichment in Dy_s (red) or St_s (blue) or no significant direction (black), respectively. **h**. PCoA on the Dy_s and St_s based on Bray-Curtis distance. Enterotype information was shown in Supplementary Figure 2.

To evaluate the health condition of an independent sample, we then constructed a cross-validated random-forest model based on the vagino-cervical microbiome of the two groups (Figure 2f). 6 bacterial species included in the classifier, *Gardnerella vaginalis, Ureaplasma* unclassified and *Prevotella bivia* were significantly enriched in the dysbiosis group (Figure 2g). We therefore classified samples using this model. In total, 244 dysbiosis samples (Dy_s) and 166 stable samples (St_s) were classified in this cohort. To be expected, St_s were almost dominated by *L. crispatus, L. iners* and *L. jensenii* (Figure 2h, Supplementary Figure 4). The type transitions of samples within subjects displayed a high level of stability longitudinally, and showed no drastically transition from Dy_s to St_s during and after probiotics supplementation compared to their baselines (Figure 3, Supplementary Figure 5). Taken together, these findings point out that women consumption of the probiotics results no shedding in vagina and had no apparent effects on re-establishing a beneficial vagino-cervical microbiome.

**Fig. 3.**
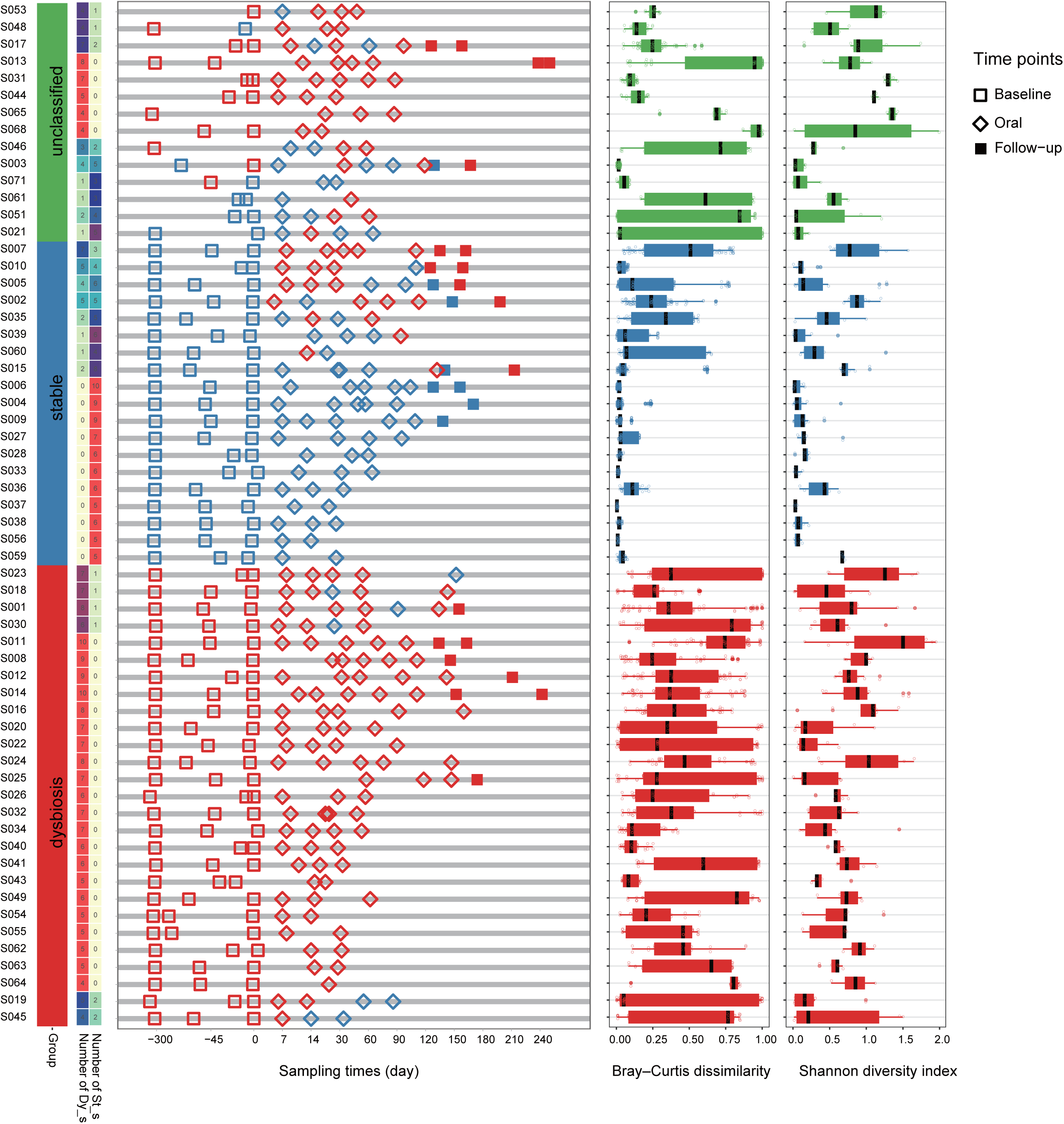
Temporal dynamics of vagino-cervical microbiome before, during and after oral probiotics. Color bar indicating dysbiosis subjects, stable subjects and unclassified subjects. Subject IDs are indicated on the left. **b**. Profiles of Dy_s or St_s samples for 60 subjects before, during and after oral probiotics. Each shape (hollow square, solid square or diamond) represents one sample in the time series. **c**. Box plot of Bray-Curtis dissimilarity between all pairs of samples within each subject. **d**. Box plot of Shannon diversity index of samples within each subject. Boxplots show median and lower/upper quartiles; whiskers show inner fences (**c** and **d**).

### Dynamics of Personalized vagino-cervical Microbiome

The vagino-cervical microbiome of 60 women were visualized by mapping temporal dynamics in community composition longitudinally (Figure 4). The microbiome composition of subjects in stable group appeared to be comparatively stable over time, and were typically dominated by *L. iners, L. crispatus* or *L. jensenii*. In these women, the slightly transitions were mostly exhibited among the different *Lactobacillus* species. The relative abundance of non-*Lactobacillus* only resides in a small space, and showed little need to improve the vagino-cervical microbial ecosystem by consumption of the probiotics. The microbiome composition of subjects in dysbiosis group changed markedly and continuously over time. However, the relative abundance of *Lactobacillus* was observed increased during and after probiotics supplementation only in 4 subjects, including *L. crispatus* in S020, *L. iners* in S030, *Lactobacillus acidophilus* in S025, and *L. iners* and *Lactobacillus* sp. 7_1_47FAA in S065 (Figure 5a-5d, Supplementary Figure 6). All the aforementioned *Lactobacillus* were present as the endogenous bacteria from the baseline period except *L. acidophilus* (Supplementary Figure 6). These results suggested that endogenous vaginal Lactobacilli could increase after the oral probiotics

**Fig. 4.**
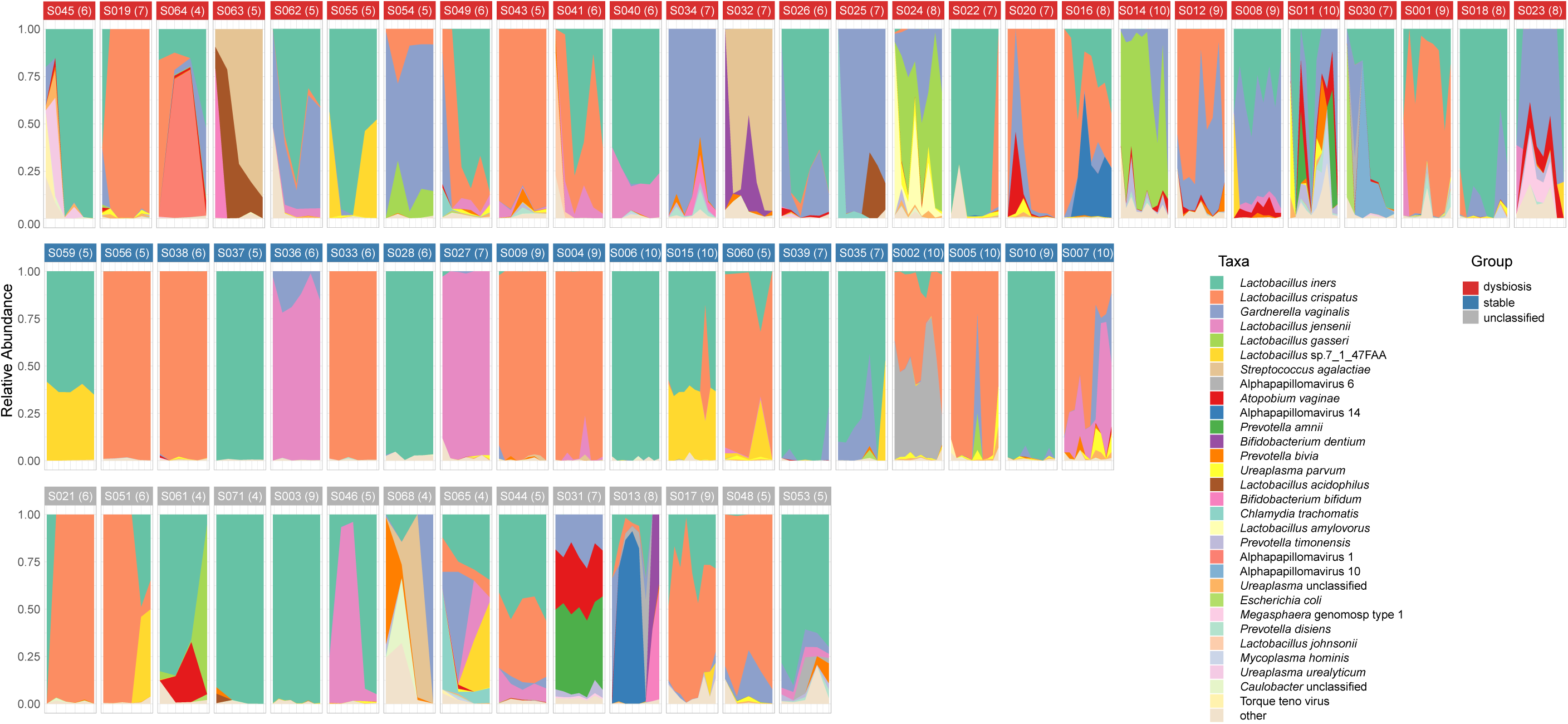
Dynamics of personalized vagino-cervical microbiome. eatmaps of the main taxa at species levels in 60 subjects is shown. Dysbiosis group, stable group and unclassified subjects present in three lines.

**Fig. 5.**
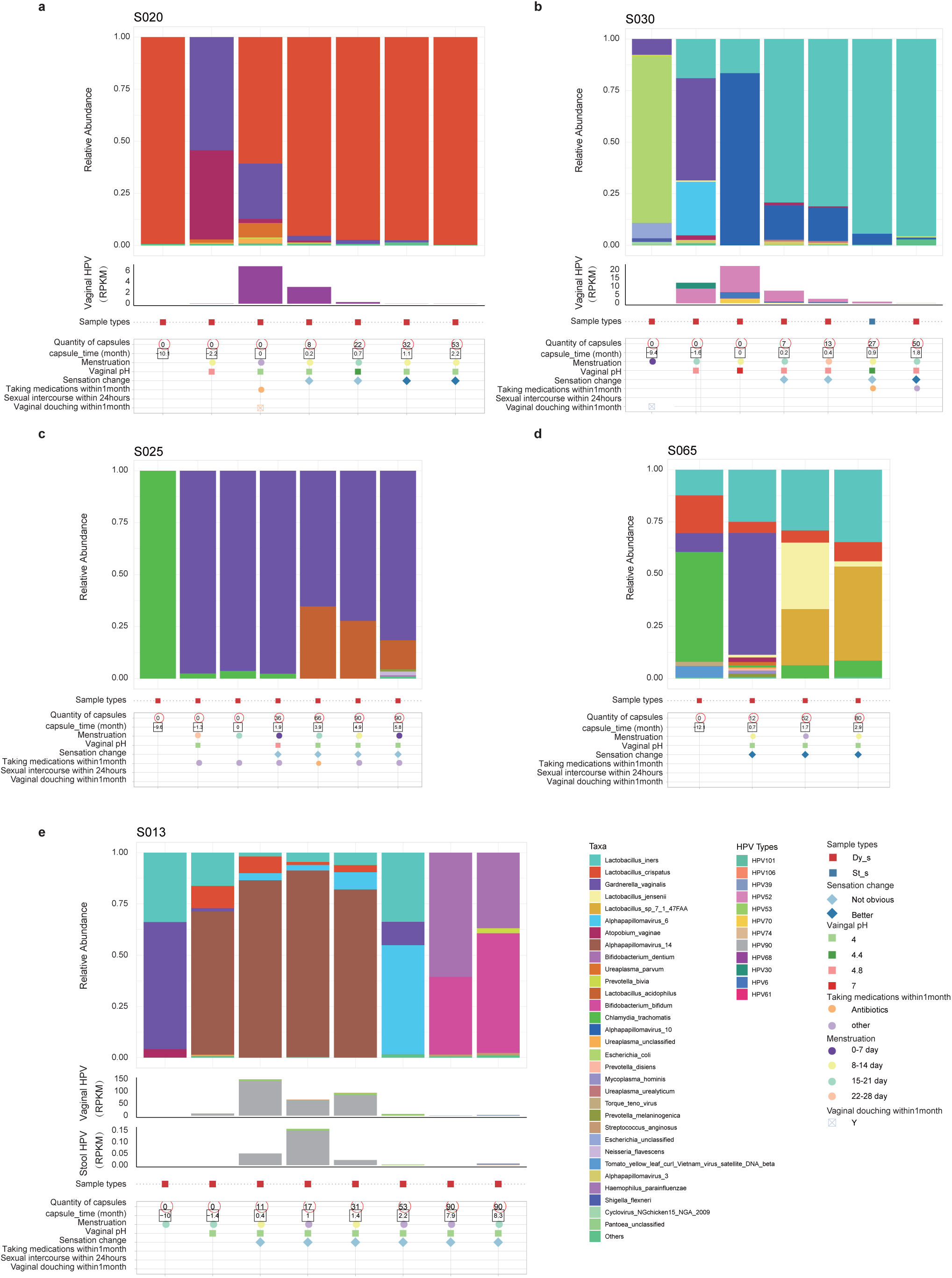
Vagino-cervical microbiome in five selected subjects. The microbial composition in each vagino-cervical sample at the species level according to MetaPhlAn2 is shown in the top. Vaginal and stool HPV types below the bar graphs were identified by HPViewer. RPKM is the abbreviation of “Reads Per Kilobase per Million reads”. Samples types including Dy_s (red) and St_s (blue). Other characteristics of subjects including quantity of capsules, capsule time, menstruation, vaginal pH, sensation changes, medical information, sexual intercourse and vaginal douching is shown in the bottom of the table.

Subjects S020, S030and S013 were detected to be infected with HPV in the baseline, but gradually be cleared away during and after their probiotic supplementation (Figure 5a, 5b, 5e). Interestingly, with the clearance away of the HPV, *Bifidobacterium* including *B. bifidum* and *B. dentium* were harboured as the dominated genus in subject S013 (Figure 5e). HPV infections were also detected in fecal samples of this subject, with a similar trend of vaginal samples in the same individual (Figure 5e). These results suggested that supplementation of these two probiotics may had some effects on HPV clearance. *Streptococcus agalactiae* (Group B *Streptococcus*), a bacterium responsible for neonatal sepsis and recently reported in placenta ^21^, could be detected in 16.7% of the subjects. But the rate of vaginal *S. agalactiae* colonization did not differ significantly between baseline and the probiotics period (*P* = 0.98, Supplementary Figure 7), consistent with colonization effects in pregnancy^19^.

## DISCUSSION

In this study, we provided metagenomic data for the first time following oral probiotics supplementation. Although some volunteers showed the Lactobacilli probiotic strains in the fecal samples, there was no increase in the probiotic strains in the vaginal or oral sites, suggesting that *L. rhamnosus* GR-1 and *L. reuteri* GR-14 were not translocated from the gut to the vagina^22^. PCR evidence of vaginal colonization has been reported for these strains for individuals with BV, and our metagenomic data raise the possibility that endogenous vaginal Lactobacilli (*L. crispatus, L. iners, etc*.) have been promoted by the oral probiotics through immunological or metabolic modulation^23^. We present the efficacy results from a comprehensive view of dysbiosis in the vagino-cervical microbiome. In volunteers with a Lactobacilli-dominated vagino-cervical microbiome, the microbiome is largely unchanged over one year, whether during probiotic intake or not. The dysbiosis group have a more diverse vagino-cervical microbiome and a less diverse fecal microbiome, but pH and microbiome dynamics varied between individuals. Without better ways of minimizing the individual dynamics, a much larger cohort would be needed to further analyze and predict the effects of probiotics supplementation.

It remains possible that *L. crispatus* could be more effective as an oral probiotic for the vagino-cervical microbiome. Other factors such as hormonal dynamics^24^, seasonal changes^25^ may also have influenced our study. Recent studies of vaginal microbial transplant (VMT) and treatment of BV using *L. crispatus* have all used a more direct topical application after standard metronidazole treatment^26,27^. Yet, oral probiotics are more readily consumed in a subclinical setting, and may be more acceptible for pregnant women with a risk for preterm birth.

### Online content

Any methods, additional references, Nature Research reporting summaries, source data, statements of data availability and associated accession codes are available at https://db.cngb.org/search/project/CNP0001123.

## Acknowledgements

This research was supported by the Shenzhen Municipal Government of China (JCYJ20170817145523036). The authors are very grateful to colleagues at BGI-Shenzhen for sample collection, and discussions, and China National Genebank (CNGB) Shenzhen for DNA extraction, library construction, sequencing.

## Author contributions

H.J. and C.C. conceived and organized this study. C.C., L.H., L.S., and X.Z. performed the sample collection and questionnaire collection. L.H., C.C., L.T, and Z.J. performed the bioinformatic analyses, H.J., C.C. and X.Z. wrote the manuscript. All authors contributed to data and texts in this manuscript.

## Methods

### Cohort demographics

With the baseline for the vagino-cervical microbiome studied from May 2017 and Feb. 2018, we started the metagenomic study for oral probiotics supplementation over the course of 3 months, followed by a two-month wash out period. The commercial probiotic capsules containing *Lactobacillus rhamnosus* GR-1 and *Lactobacillus reuteri* GR-14, and each capsule at 2.5 billion colony forming units (CFUs). The study was approved by the Institutional Review Boards at BGI-Shenzhen (IRB approval numbers 17244). 60 healthy women aged from 23 to 61 were recruited in Shenzhen, China (Supplementary Table 1). Exclusion criteria included: (i) Pregnant women, (ii) consumption of probiotics or antibiotics in any form within one month prior to participation. All participants provided written informed consent at enrolment, and then received a first online questionnaire covering comprehensive demographic characteristics (Supplementary Table 1). The study design consisted of three phases, baseline (10 months), probiotic intervention (consumed 90 capsules of probiotics) and follow-up (2 months). Samples were collected three times during the baseline phase (B2-B0). Time point B2 was about 10 months before probiotic intervention, and B1 was about 1.5 months before probiotic intervention. B0 was the most recent time point, participants were instructed to collect samples after menses period, then began to received probiotic capsules. During the intervention phase, each participant was assigned 90 capsules of probiotics and instructed to take one capsule daily. Samples were scheduled 7 (O1), 14 (O2) days after intervention, then monthly after menses period throughout the rest of the intervention (O3, O4, O5). After intervention, two follow-up visits were scheduled monthly after menses period (W1, W2). Vaginal samples were collected at each time point using a home collection kit. Two vaginal swabs were requested, the swab head of one was put into tube with storage reagent (ref), the other one was brushed on the pH test strips. Other three different kinds of samples (buccal mucosa samples, tongue coat samples and fecal samples) were also collected by self-sampling at all time points except B2, B1. Participants were also requested to fill in an online questionnaire at each time point. The information of questionnaire including vaginal PH value, sampling time, menstruation, sexual activity. Samples belonged to probiotic intervention were removed when the participant’s average capsule of probiotics was less than 0.5 a day. Throughout the entire study 1334 samples including 322 tongue coat samples, 263 buccal mucosa samples, 436 vaginal samples and 313 fecal samples were collected.

### DNA extraction and metagenomic shotgun sequencing

DNA extraction of all samples from four body sites was performed as described^28^. Metagenomic shotgun sequencing was performed on the BGISEQ-500 platform (100bp of paired-end reads) ^29–32^. The sequencing reads of stool samples were quality-controlled using Overall Accuracy (OA) control strategy (https://github.com/Scelta/OAFilter), and then aligned to hg19 to remove human reads using SOAP2.22 (SOAPaligner/soap2, RRID:SCR_005503) as described previously^29^. Stringent condition for removal of host sequences was used for tongue coat samples, buccal mucosa samples and vaginal samples^32^, through alignment to the hg19, hg38 and YH reference by DeconSeq ^33^(version 0.4.33) and SNAP^34^. Taxonomic assignment of the high-quality metagenomic shotgun data of samples from four body sites were performed using MetaPhlAn2^35^ version 2.7.0 with database v20. HPViewer with the default parameters was used to detect genotyping of HPV in the high-quality metagenomic sequencing data of samples^36^.

### Statistical analysis

Alpha diversity and beta diversity were calculated on species relative abundances using Shannon-Wiener index and Bray-Curtis dissimilarity, respectively. Kruskal-Wallis test was used to make temporal dynamics comparisons among different time points, including species relative abundance of oral probiotics, Shannon-Wiener index, Bray-Curtis dissimilarity, vaginal pH and relative abundance of *Lactobacillus* genera. Wilcoxon rank sum (Mann-Whitney U) and Wilcoxon signed-rank test were used to make comparisons between two groups or each of two time points. The statistical significance was with a p value threshold of 0.05 and a false discovery rate (FDR) threshold of q < 0.05.

To build a predictive model to identify microbial dysbiosis, the species relative abundances in the baseline samples were calculated with the training set with 500 trees in the random Forest package (version 4.6-14). Five-fold cross-validation was performed five times. The cross-validation error curves from the five trials were averaged, and the minimum error in the averaged curve plus the standard deviation at that point were used as the cutoff for acceptable error. From the sets of species with a classification error less than the cutoff, the set with the smallest number of species was chosen as the optimal set, as in previous methods on the vagino-uterine microbiome^28^. Relative abundances of species in all 410 vaginal samples were used to determine the optimal community types of the vagino-uterine microbiome according to hierarchical clustering based on the Jensen-Shannon distances and Ward linkage. And more statistical details were described in the results and denoted in figure legends, including sample summary, distribution, the statistical method and the statistical test used and significance.

## Data availability

Metagenomic shotgun sequencing data for all samples have been deposited to the (CNGB) database under the accession code CNP0001123.

