## Supplementary figures for "Personalized vagino-cervical microbiome dynamics after oral probiotics"

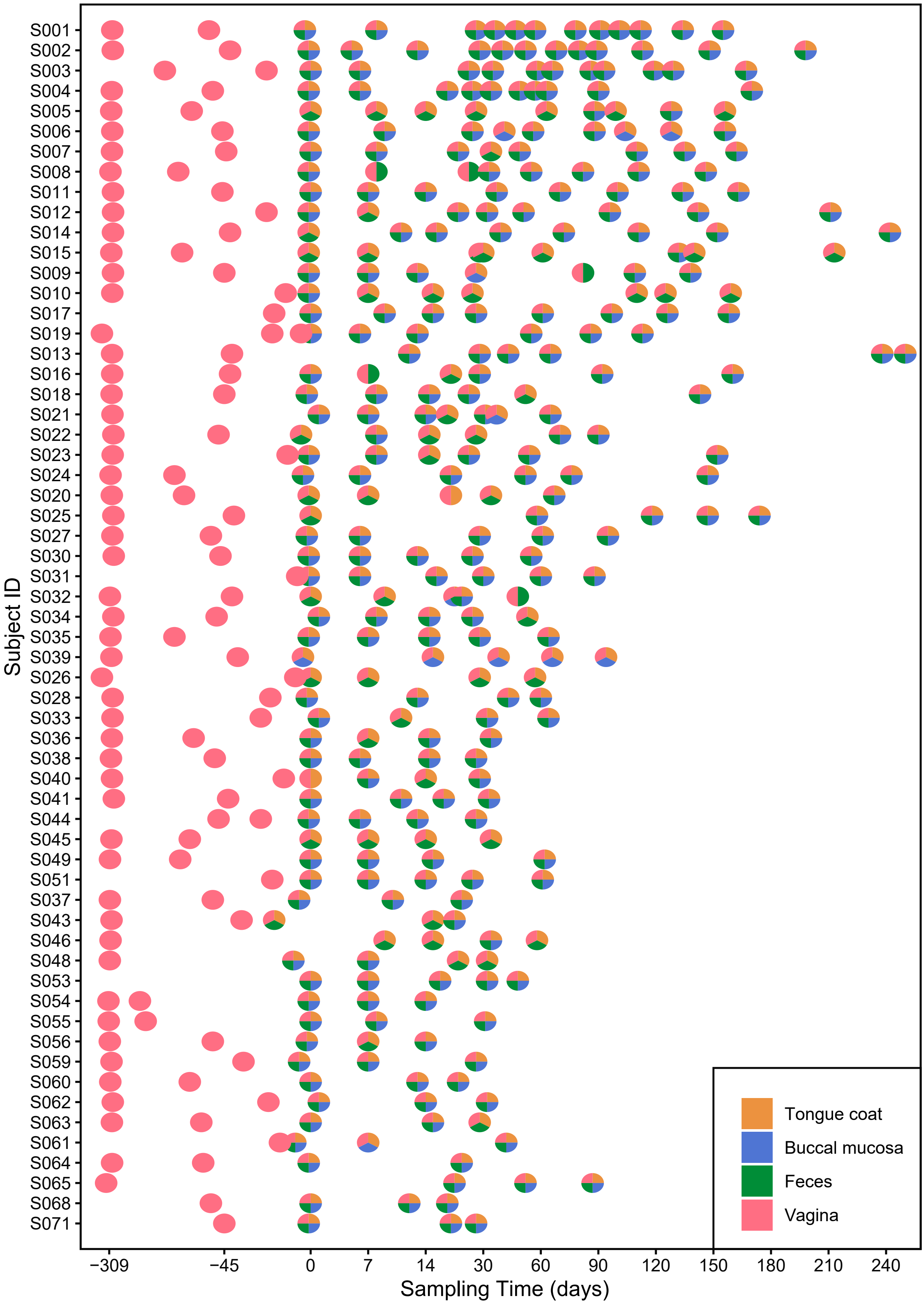


**Supplementary Figure 1. Summary of subjects sampled along the time axis in our cohort.**

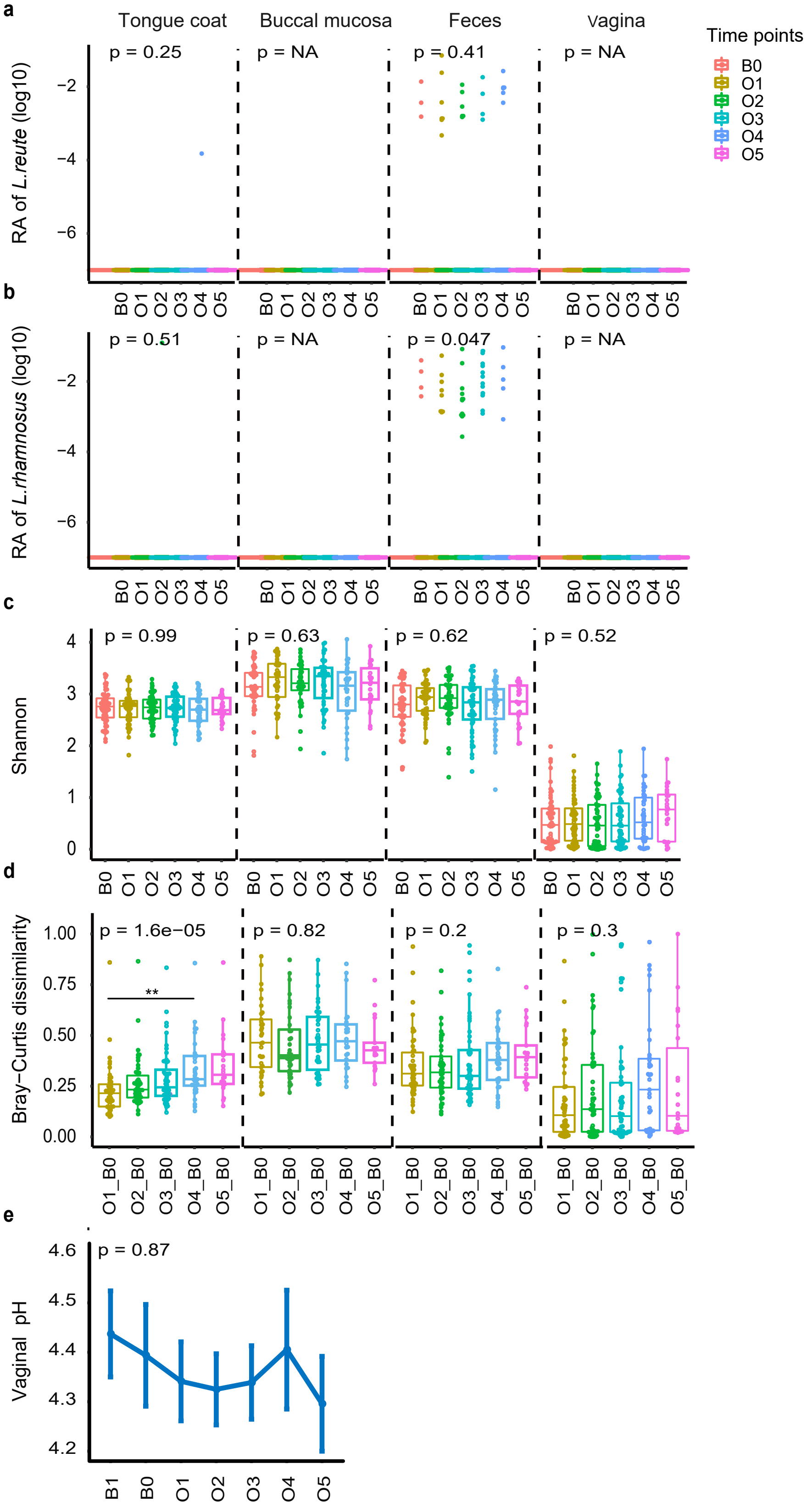


**Supplementary Figure 2. Lack of oral probiotic colonization in the microbiome from the multiple body sites before and during oral probiotics.** Relative abundance (RA) distributions of *L. reuteri* RC-14 (**a**) and *Lactobacillus rhamnosus* GR-1 (**b**). **c.** Shannon diversity index. **d.** Bray-Curtis dissimilarity between B0 and other sampling points within individuals. **e.** Vaginal pH. Boxplots show median and lower/upper quartiles, and whiskers show inner fences (**a-d**). Kruskal-Wallis test is used to conduct temporal dynamics comparisons (**a-e**).

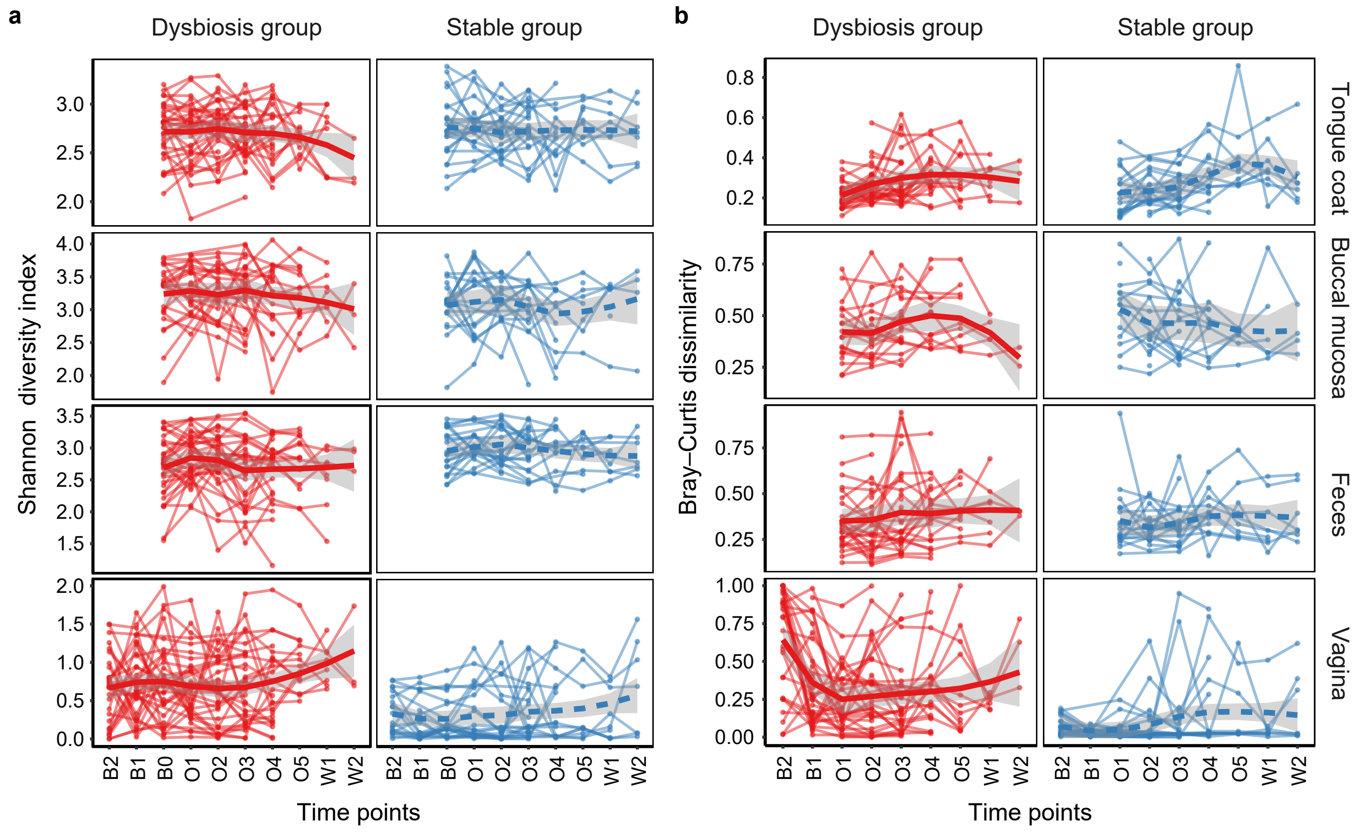


**Supplementary Figure 3. Temporal shifts in the microbiome from the multiple body sites in dysbiosis and stable groups. a.** Shannon diversity index. **b.** Bray-Curtis dissimilarity between B0 and other time points within individuals.

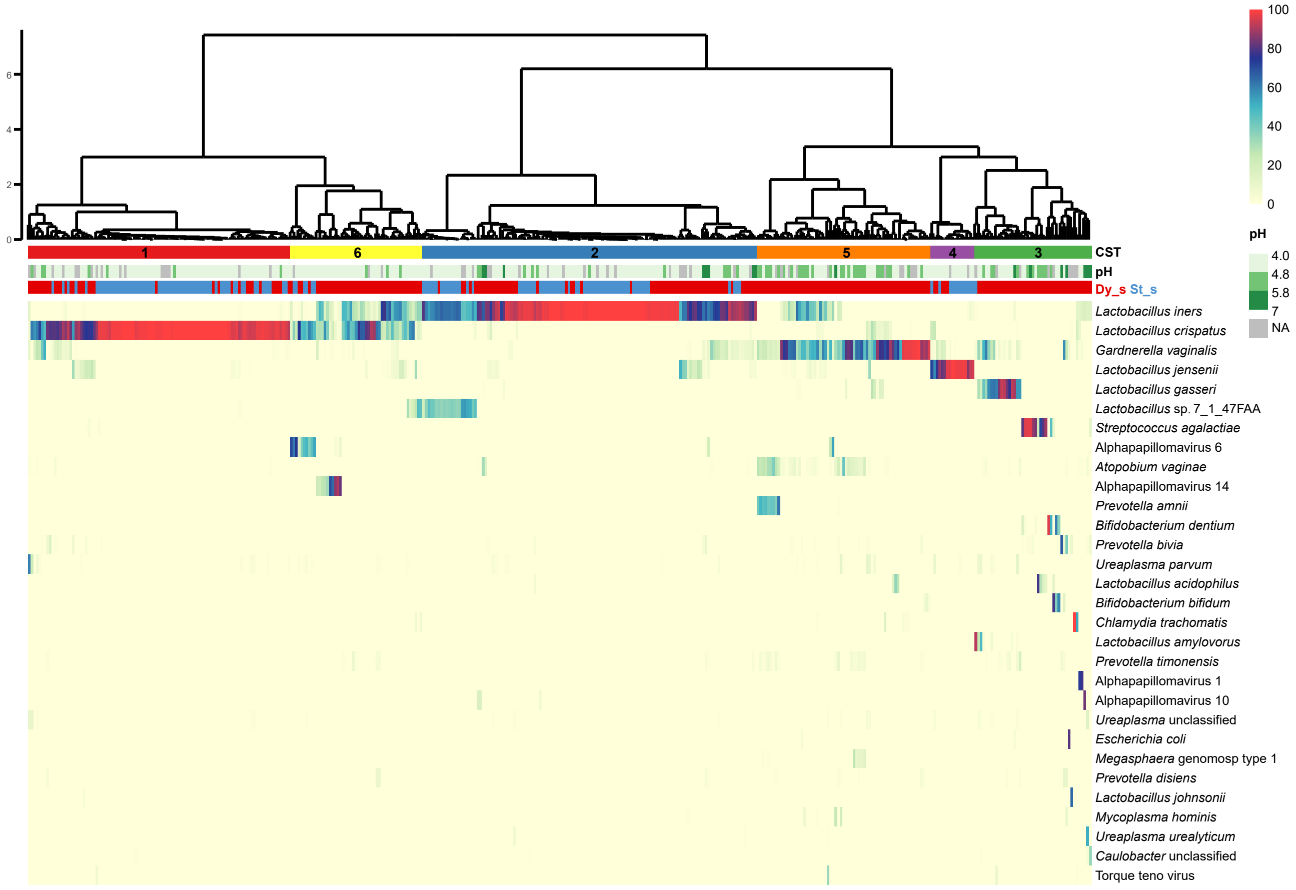


**Supplementary Figure 4. Heatmap of the fractional abundance of the 30 most abundant species in the vagino-cervical microbiome of 60 women sampled longitudinally during before, during and after oral probiotics.** The samples are hierarchically clustered (R base hcluster function with Jensen-Shannon distances and Ward linkage, and identified six community state types (CST). CST 1, 2, 4, 5 are characterized by dominant *Lactobacillus crispatus*, *Lactobacillus iners*, *Lactobacillus jensenii* and *Gardnerella vaginalis*, respectively. In CST 6, *Lactobacillus crispatus*, *Lactobacillus iners* and Alphapapillomavirus constituted notable fraction of the microbiome. CST 3 is dominated by *Lactobacillus gasseri* or *Streptococcus agalactiae* or *Bifidobacterium dentium* or some other species. Vaginal pH and samples types including Dy_s (red) and St_s (blue) are indicated by the bars, respectively.

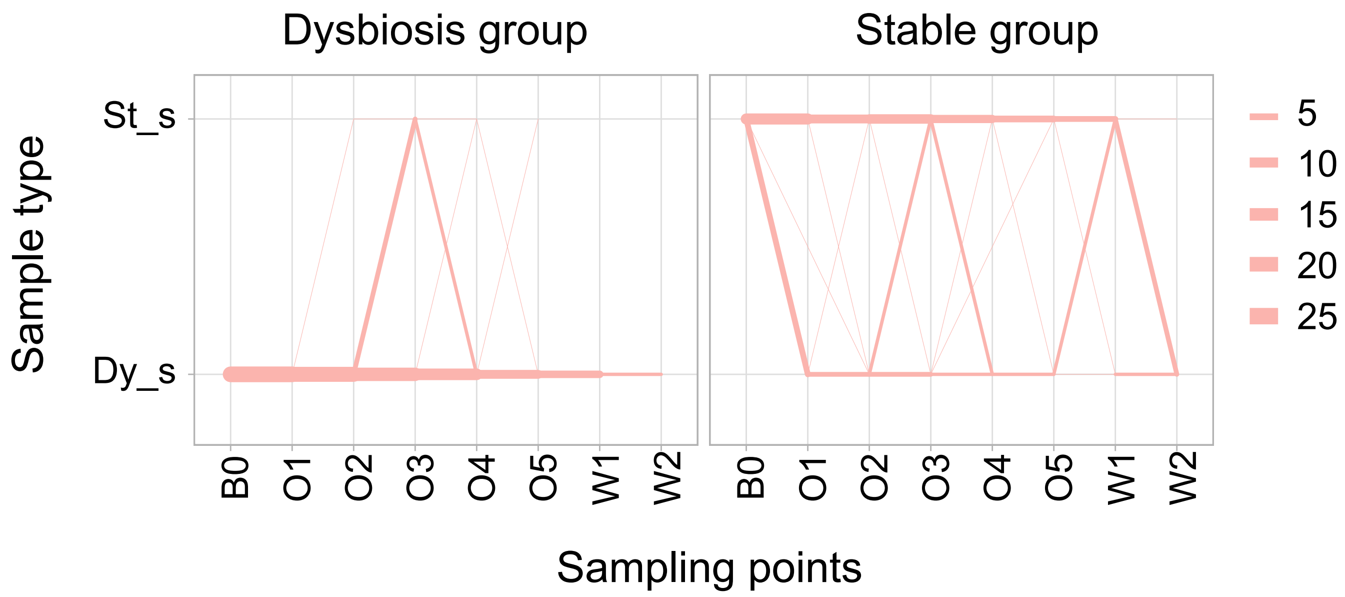


**Supplementary Figure 5. Transitions of sample types in Dysbiosis and stable subjects, respectively.**

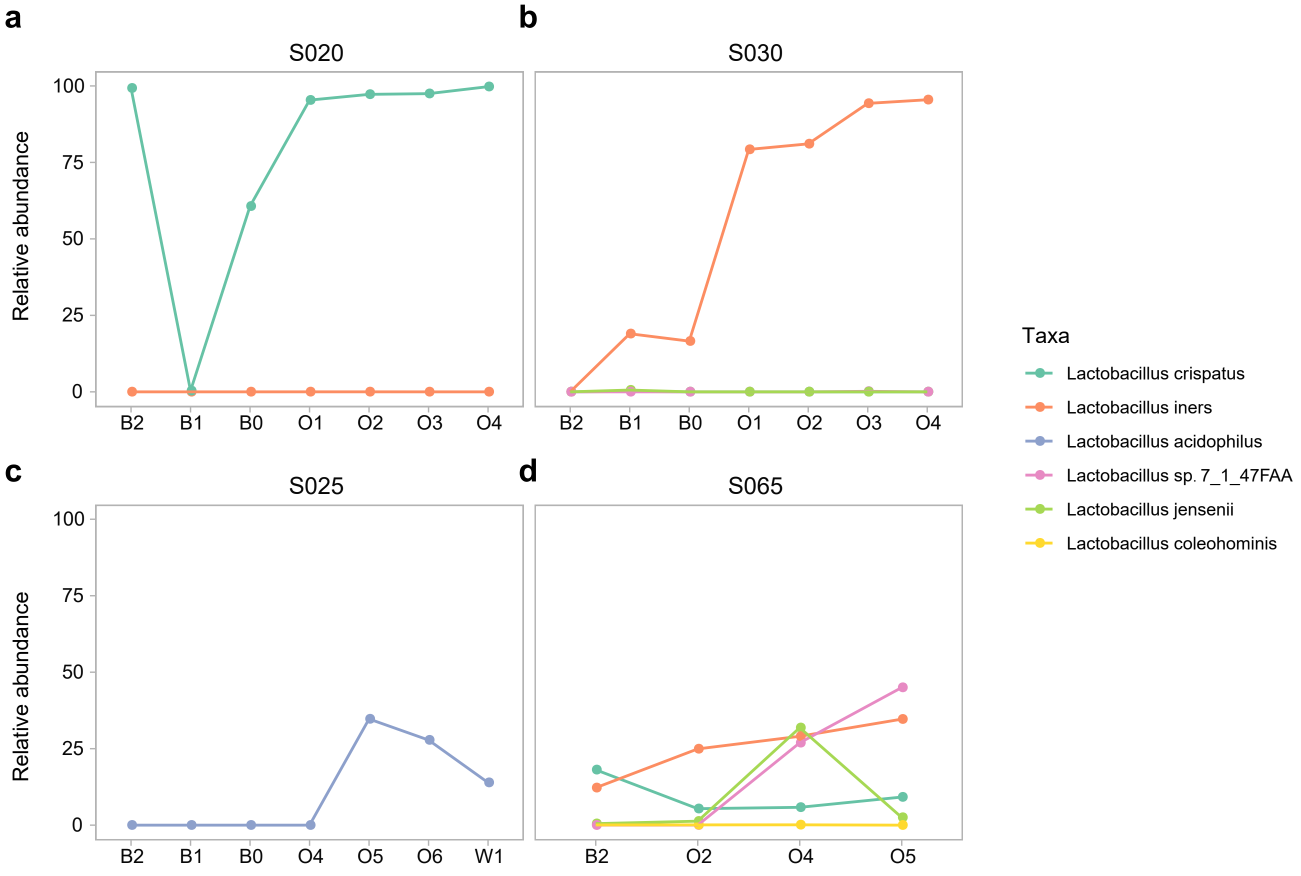


**Supplementary Figure 6. Relative abundance of *Lactobacillus* in the vagino-cervical microbiome of four *Lactobacillus*-increasing subjects.**

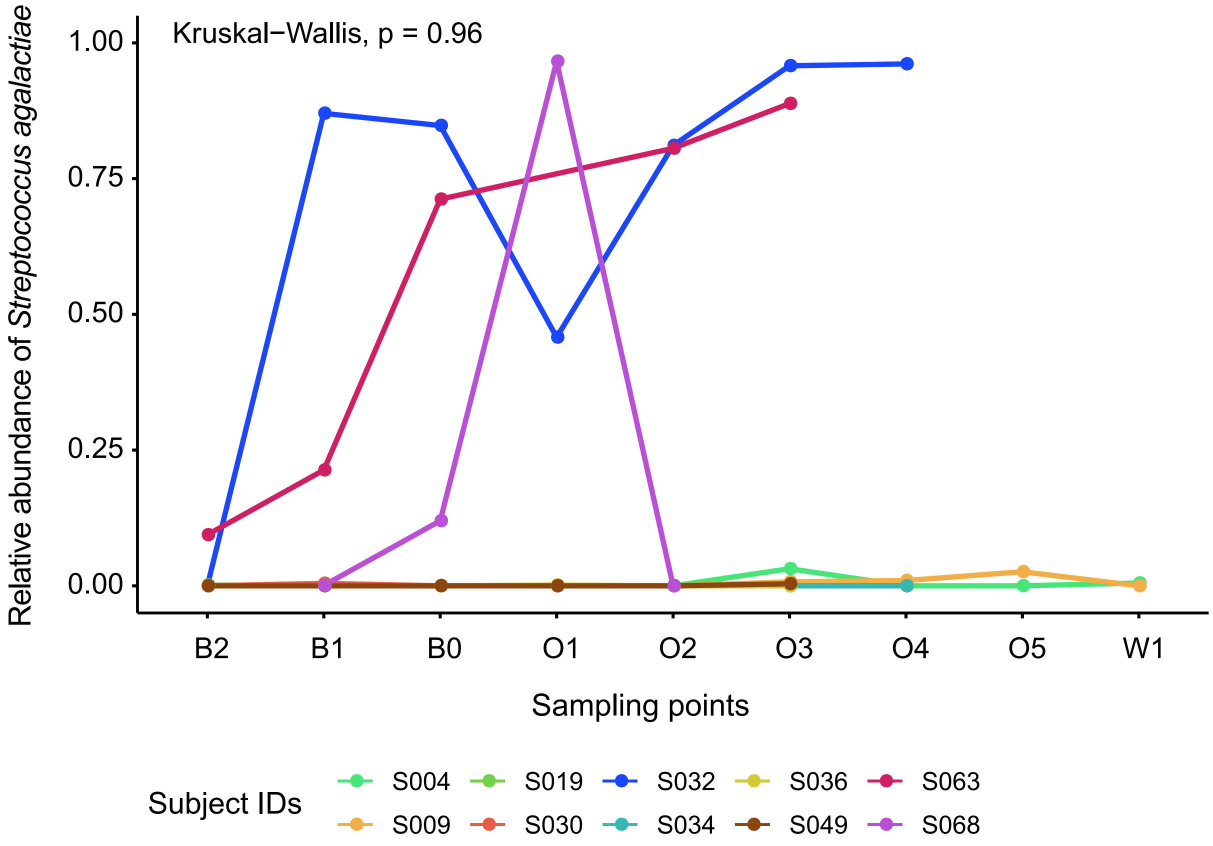


**Supplementary Figure 7. Relative abundance of *Streptococcus agalactiae* in the vagino-cervical microbiome before, during and after oral probiotics.** *S. agalactiae* is detected in only 10 subjects. Kruskal-Wallis test are used to conduct temporal dynamics comparisons, and no significant changes within subjects during and after oral probiotics compared to their baseline (P=0.96).
